# Development of shell field populations in gastropods

**DOI:** 10.1101/2024.05.16.592602

**Authors:** Supanat Phuangphong, Hiroki Yoshikawa, Yune Kojima, Hiroshi Wada, Yoshiaki Morino

## Abstract

The embryonic shell field of mollusks first appears during gastrulation of the dorsal ectoderm and subsequently develops into the shell-secreting mantle in adult animals. Although several lines of evidence have revealed that this shell field lineage is exclusively derived from the second quartet (2q) of the 16-cell embryos, it is generally believed that the establishment of the shell field fate would be accomplished only after receiving inductive signals from the invaginated endoderm. Despite being accepted as a comprehensive model for molluskan shell field specification, the validity of this induction hypothesis remains questionable owing to the lack of clear experimental evidence and contradictory results. Here, we attempted to re-investigate the inductive role of the endoderm in shell field fate establishment in the limpet *Nipponacmea fuscoviridis* by experimentally disrupting cell-cell contacts between cell lineages after the 16-cell stage. First, we characterized the shell field cell population by performing two-color *in situ* hybridization. We characterized at least three cell populations in the developing shell field. Using single-cell transcriptome analysis, we identified several specific effector genes for each population, as well as transcription factor genes. Differentiation of each shell field population was inspected in 2q blastomeres isolated from other cells of the 16-cell embryos. Despite the absence of any interlineage interactions (including ectoderm-endoderm contacts), the expression of marker genes for each shell field population was observed in the isolated 2q fragments. In addition, the expression of several shell field genes was detected in embryos in which cytokinesis was blocked at the 16-cell stage. We concluded that the early process of shell field differentiation in the 2q lineage occurs mostly independently of the interactions with other lineages.

## Introduction

One of the hallmark characteristics of the phylum Mollusca is the presence of dorsal shells, which has evolved into various forms in distinct molluskan groups. As a first step toward understanding the evolutionary history of molluskan shells as a novel characteristic, we attempted to elucidate the developmental mechanism of the shell. Like other spiralians, most mollusks display spiral cleavage with early cell-fate segregation among the blastomeres along the animal-vegetal axis; thus, the major cell lineages of the embryo usually appear just after a few rounds of cleavage (1). The major blastomere lineages that contribute to the shell field tissues vary among different mollusk taxa; however, most generally emerge from second quartet (2q) derivatives. Examples of this are emergence from 2a–d in the gastropods *Patella vulgata*, *Tritia* (*Ilyanassa*) *obsoleta*, and *Crepidula fornicate*, as well as emergence from the 2d in the bivalves *Mytilisepta virgata* (*Septifer virgatus*) and *Unio complanata* and in the scaphopod *Dentalium* (2–7) The specification of each quartet lineage may be achieved by interactions among the spiralian-specific homeobox gene SPILE(8, 9). However, subsequent processes toward the specification of shell field cells remain to be elucidated. The acquisition of the shell field fate may be achieved later by cell-cell interactions between different germ layers. Raven (10) proposed the first model of shell field induction regarding the role of the invaginated endoderm based on experiments in the gastropod *Lymnaea stagnalis*. During gastrulation, endodermal cells invaginate at the vegetal pole and extend deeper inside the blastocoel until the tip of the invaginated tissue touches the overlying sheet of the dorsal ectoderm. After ectoderm– endoderm contact occurs, dorsal epidermal cells begin to differentiate into more elongated shell field cells. The elongated dorsal ectoderm undergoes invagination before it soon undergoes evagination. These sequential morphogenetic events are referred to as shell field invagination and evagination, respectively (10–12). The evaginated shell field then expands and covers the dorsal part of the developing animal with a secreted shell matrix (12). Raven (10) found that if physical contact between the dorsal ectoderm and invaginated endoderm was interrupted by treating embryos with LiCl, the dorsal ectoderm failed to differentiate into a shell field. This suggests that the dorsal ectoderm probably acquired the shell field fate only after it came into contact with the underlying endodermal cells and received an inductive signal.

Since the proposal of this shell field induction model, consistent results have been revealed in several studies, although the specific molecule acting as an inducer has not yet been uncovered (7, 12, 13). However, contradictory results and problematic aspects of this model have been revealed. Collier and McCann-Collier (14) found that the shell field of the gastropod *T. obsoleta* had already formed when the archenteron had initially developed. Moreover, not only the shell field cells but also the other epithelial cells appeared to contact the underlying 4D macromere. Therefore, they suggested that induction of the shell field by the endoderm was unlikely in this species.

These contrasting conclusions regarding the acquisition of shell field fate have posed difficulties in creating a comprehensive picture of molluskan shell field development and a complete understanding of the embryonic development of this animal group. One of the critical reasons for this inconsistency is the lack of proper shell development indices. Most of the 20^th^ century studies have evaluated shell development by observing birefringence using polarized light. Given that gastropod embryos also secrete the matrix in the operculum and statocyst (15, 16), we cannot exclude the possibility that matrix secretion in the deformed larvae reflects the differentiation of the operculum or statocyst instead of the shell field cells (17).

In addition, the shell field is not a uniform group of cells; rather, it is composed of multiple cell types. Although the function of each cell layer in the larval shell field remains unclear, the adult mantle is composed of multiple cell types, each of which has a different function in the formation of the adult shell (18–20).

In this study, we reinvestigated the role of cell-cell interactions in the shell field fate establishment of the limpet *Nipponacmea fuscoviridis* by attempting to interrupt cell-cell contact. We attempted to eliminate all inter-lineage interactions between the 2q blastomeres and others by isolating the 2q blastomeres for independent development. Six shell field-associated genes, including three transcription factor (TF) genes (*Hox1*, *Mist* and *Hox4*), one signaling molecule (*dpp*), and two effector genes (Tyrosinase [*Tyr*] and chitin synthase 1 [*CS1*]) were used as markers for shell field differentiation. These molecular markers allowed us to inspect differentiated shell field cells without relying on morphological clues. Here, we found that, despite the interruption of possible interlineage communication, the signals of all marker genes were still detected in the cell clusters derived from the isolated 2q blastomeres. Our results are not consistent with the proposed model of shell field induction and indicate that the expression of early shell field genes is activated in a cell-autonomous manner in 2q lineage cells.

## Materials and Methods

### Animal acquisition, artificial fertilization, and drug treatment

Sexually mature samples of the limpet *N. fuscoviridis* were collected from Hiraiso (Ibaraki, Japan). *In vitro* fertilization was performed as previously described (15). After fertilization, embryos were incubated at 22°C in artificial seawater (ASW). For inhibition of cytokinesis, 16-cell limpet embryos were treated with 5 μg/mL cytochalasin B (Fujifilm) in ASW until nine hours post fertilization (hpf), and then fixed.

### Whole-mount *in situ* hybridization

The primers used for gene isolation are listed in Table S1. Digoxigenin (DIG)- or fluorescein-labeled probe synthesis was performed following the protocols described by Yamakawa et al (21). *In situ* hybridization was conducted according to previously described protocols (9). For two-color *in situ* hybridization, fluorescein- and DIG-labeled RNA were added to the hybridization buffer. Anti-DIG-alkaline phosphatase (AP) (Roche) and anti-fluorescein-AP (Roche) were used for the antibody reaction. Nitro blue tetrazolium/5-bromo-4-chloro-3-indolyl phosphate (Roche) and Fast Red (Sigma-Aldrich) were used to visualize the probe distribution. After staining with the first probe, the sample was treated with 0.1 M glycine-HCl (pH 2.2) for 15 min at room temperature to inactivate the antibody, followed by staining with the next probe.

### Single-cell transcriptome analysis

#### Sample preparation, library construction, and sequencing

Early trochophore larvae (8 hpf) were collected in a 15-mL tube and washed three times with Mg/Ca-free ASW. They were then rocked in a 2% trypsin solution dissolved in Mg/Ca-free ASW for 45 min at room temperature and gently pipetted every 10 min. The dissociated larvae were then transferred to a Petri dish and pipetted repeatedly into a dissociation buffer (1.0 M glycine, 25 mM ethylenediaminetetraacetic acid, pH 8.0) on ice for 10 min to dissociate into single cells. The cells were filtered using 41-µm nylon mesh and washed three times with Mg/Ca-free ASW, and centrifuged for 5 min at 3000 × g. Fluorescein diacetate and propidium iodide were used to measure cell viability and the viable cell ratio was confirmed to be greater than 90%. Single-cell library construction was conducted in a 10X chromium system using Chromium Next GEM Single Cell 3’ Reagent Kits v3.1. Library sequencing was performed by Macrogen using a HiSeq X Ten platform.

#### Analysis of single-cell RNA sequencing (RNA-seq) data

The *de novo* reference sequences were generated from the transcriptome data. We extracted total RNA from nine developmental stages and six adult organs and conducted RNA-seq (detailed information is described in Text S1). The obtained fastq sequences were filtered and trimmed using Trimmomatic (v.0.37) (22) and assembled using Trinity (v2.8.4) (23). To reduce redundancy, we used tr2aacds2.pl in the evidential genes (24) with the-MINSCD=100 option. The quality of the generated assembly was assessed using BUSCO (v4.0.6) (25) with the metazoan odb10. These assembly were used as references for subsequent analyses.

We used CellRanger (v6.12) (26) “mkref” and “count” function with the default settings. The obtained cell count was analyzed in R v4.1.3(27) using the Seurat v4.1.1 package (28). Cells with fewer than 200 or more than 5000 expressed genes were excluded. Counts were normalized using NormalizeData (normalization.method = “LogNormalize”, scale.factor = 10000) and highly variable genes were detected (FindVariableFeatures; selection.method = “vst”, nfeatures = 2000). The counts were then log scaled (ScaleData function). A principal component analysis (RunPCA) was conducted using 2000 highly variable genes. The top 20 principal components were used in subsequent analyses. We constructed a nearest-neighbor graph using FindNeighbors, and the clusters were determined using the FindClusters function (resolution = 0.5). We visualized transcriptomes with UMAP (Uniform Manifold Approximation and Projection) using the RunUMAP function. The marker genes for each cluster were identified using the FindAllMarkers function (only pos = TRUE, min.pct = 0.25, and logfc.threshold = 0.25). The top 50 marker genes in each cluster were annotated using BLAST against UniProt and gene models of oyster *Crassostrea gigas* (29) (e-value threshold =0.0001) (Table S2). The annotations for each cluster other than the shell field are provided in Text S2 and Figures S2-3. Output data of CellRanger and reference assembly sequences are available in the GEA database (E-GEAD-737).

#### Blastomere separation

All culture plates containing the isolated cells were coated with horse serum to prevent cell adhesion to the plate walls. To remove the egg-coating materials that impede the blastomere separation processes, four-cell embryos were briefly suspended in a dissociating solution (0.5% thioglycolate and 0.25% pronase in calcium-free seawater (CFSW)), followed by gentle shaking on a plate shaker. The embryos were then repeatedly washed with CFSW before being transferred to a new plate filled with clean CFSW. To accurately isolate 2q cells, the isolation processes were carried out in a stepwise manner (Fig. S1). At the third cleavage, larger 1Q blastomeres were separated from smaller 1q cells using a micropipette and glass needles. After the isolated 1Q cell underwent the next round of cell division, the smaller (2q) and larger cells (2Q macromeres) were separated. Isolated 2q and 2Q cells were transferred to separate plates filled with clean ASW. These isolated blastomeres were cultured at 22°C until they were fixed at 9 hpf.

#### Phylogenetic analysis for transcription factors

Phylogenetic analysis was performed to confirm the orthology of the TFs. The amino acid sequences were aligned with TF sequences of bilaterian species using MAFFT (30) (v 7.490) with LINSI option. The conserved domain sequences of each gene were used for phylogenetic analysis (Fig. S4). The aligned sequences for phylogenetic analysis were provided in Dataset S1. Substitution models were inferred using ModelTest-NG v0.1.7 (31), and maximum likelihood trees were constructed using RAxML-NG version 1.1 (32).

## Results

### The spatiotemporal expression of the shell field genes

First, we investigated the expression dynamics of shell field genes during normal shell field development in *N. fuscoviridis*. We found that *Hox1* and *Mist* displayed similar expression dynamics both spatially and temporally (Fig. 1A–L, A’–L’). The expressions of both genes were first detected at 5 hpf in four groups of cells around both the dorsal and the ventral sides of an embryo (fig. 1A, G, A’, G’), probably in the descendant cells of all four 2q blastomeres (A, B, C and D). The expression, then, gradually became restricted to the dorsal area during 6–9 hpf (Fig. 1B–E, B’–E,’ H–K, H’–K’). The expression appeared as a semicircular stripe at 9 hpf when observed from the dorsal view (Fig. 1E, K). These dynamics of *Hox1* and *Mist* expressions are consistent with the pattern of cell movements previously observed during shell field development in *L. goshimai* (33). At 10 hpf, the expression domains of both genes were limited to the dorsal post trochal region, whereas the signals transited from the semicircular stripe into a U-shaped pattern bordering the center of the shell field (Fig. 1F, F’, L, L’).

**Figure 1.**
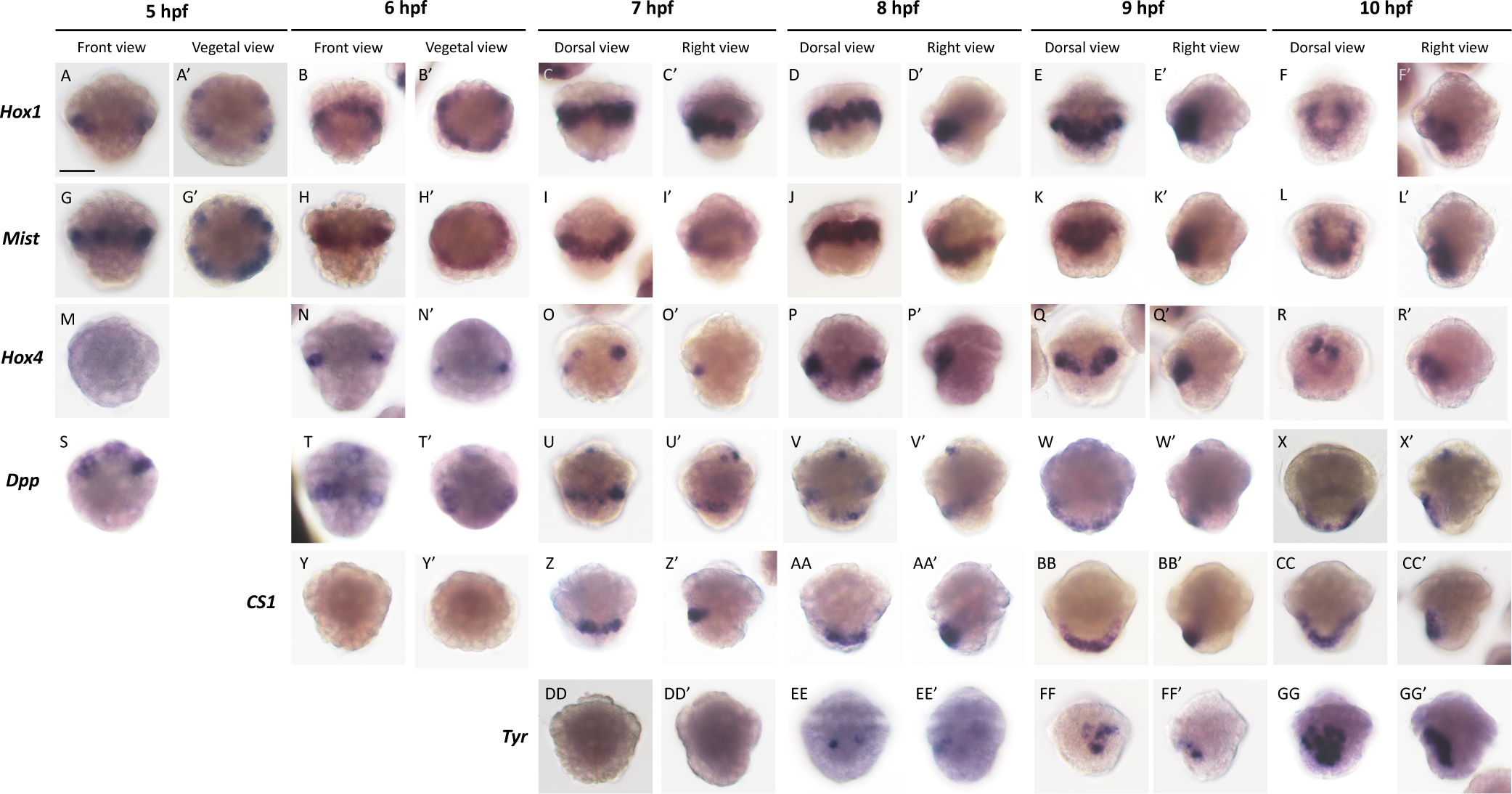
The spatiotemporal expression patterns of the shell field genes. The expression patterns of the shell field-related genes are displayed as the front or dorsal view and the vegetal or right-side view. Scale bar = 50 μm.

The expressions of Hox*4*, *dpp* and *CS1* in the shell field region were detected later than the onset of *Hox1* and *Mist* (Fig. 1M–CC, M’–CC’). Two separated centers of *Hox4* expression were initially observed at 6 hpf in two separated groups of the developing shell field cells (Fig. 1N, N’). These two groups of signals then expanded and moved toward each other on the dorsal side during 8–9 hpf until they merged into a single domain at the center of the shell field at 10 hpf (Fig. 1O–R, O’–R’). The post trochal expression of *dpp* was also detected at 6 hpf in two separate groups of cells, although expression was detected at an earlier stage in the pretrochal ectoderm (Fig. 1 S –T, T’). The *dpp* expression was detected on the vegetal side of the shell field at later stages (Fig. 1U–X, U’–X’). The expression of *CS1* first appeared at 7 hpf in a thin stripe located at the vegetal side of the shell field (Fig. 1Z, Z’). The expression of *CS1* in the later stages was similar to that of *dpp* on the vegetal side of the shell field (Fig. 1AA-CC, AA’-CC’). The spotted expression of *Tyr* was detected in the developing shell field at 8 and 9 hpf (Fig. 1EE-FF, EE’-FF’). At 10 hpf, the expression of Tyr was continuously observed in the shell field, with an increasing number of Tyr-positive cells. (Fig. 1GG,GG ’).

We found that the shell field genes appeared to be expressed in distinct groups of cells during shell field development, and to further describe the relative positions of these populations of shell field cells, we carried out two-color *in situ* hybridization in the 7- and 9-h embryos for the following gene pairs: *Mist*-*CS1*, *CS1*-*Hox1*, *Hox4*-*CS1 Tyr*-*CS1*, *Mist-Dpp*. At the 7-hpf stage, at least three distinct groups of developing shell field cells were revealed based on the expression patterns of shell field genes (Fig. 2A–F). The anterior part of the developing shell field, which was located directly posterior to the prototroch, was defined by the expression of both *Hox1* and *Mist* (Fig. 1), whereas the posterior part was composed of *CS1*-positive cells that were located immediately posterior to the *Hox*1-*Mist* positive region (Fig. 2A–D). *Hox4*-positive cells were found on the anterior sides of the shell field that were not in contact with *CS1*-positive cells, indicating that *Hox4* is expressed anteriorly and bilaterally in both sides of the *Mist* and *Hox1* expression domain. (Fig. 2E–F).

**Figure 2.**
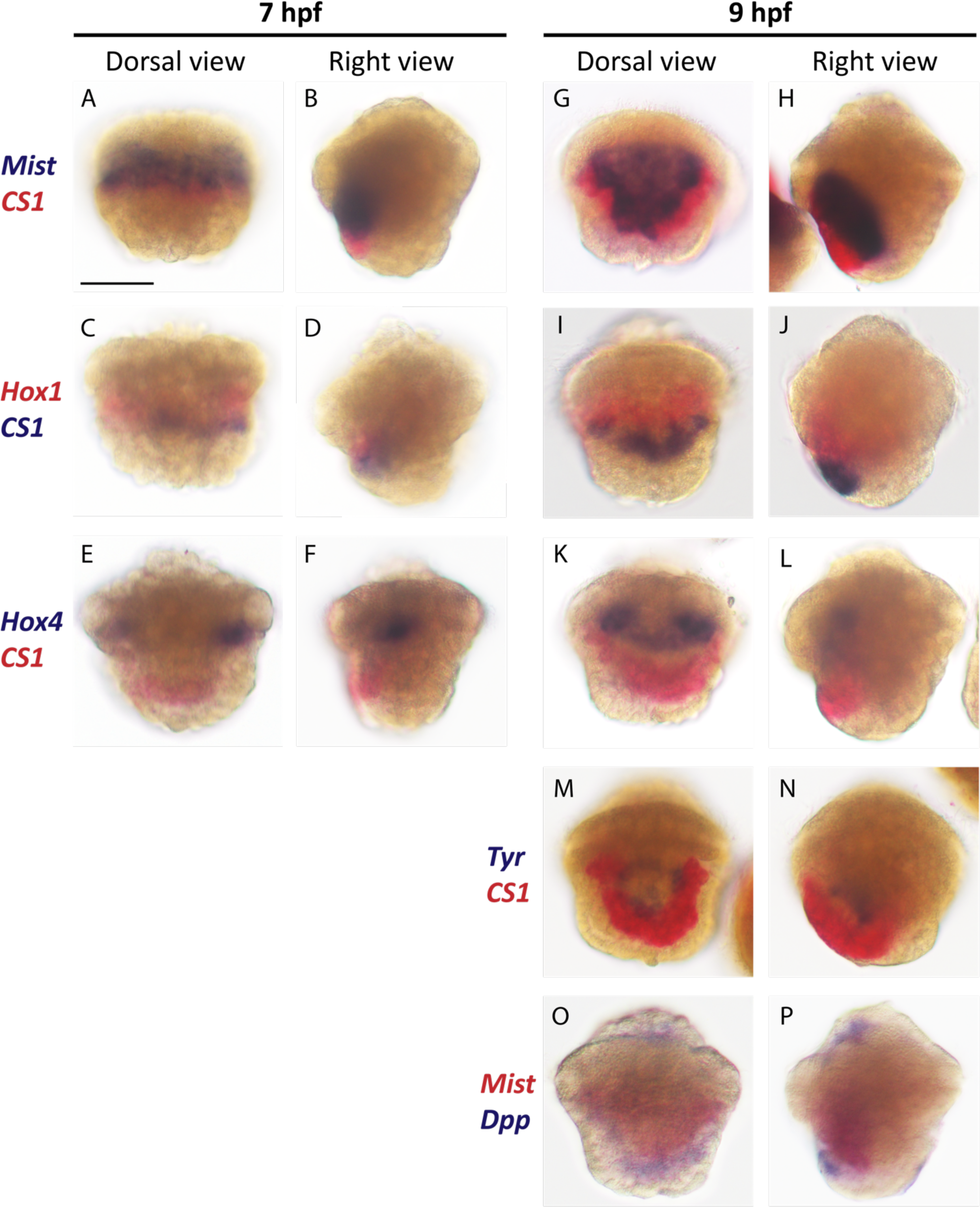
Double staining of shell field genes. The expression patterns of shell field genes are represented in both the dorsal and right -side views. Scale bar = 50 μm.

At the 9-hpf stage, *Dpp* was then expressed in the band of the cell posterior to the *Mist-* positive cells, indicating overlapping expression with *CS1* (Fig. 2O and P). The expression of *Tyr* was sparsely observed in some of the cells at the center of the shell field, which was bordered by *CS1*-positive cells on the bottom and both sides of the U-shaped pattern (Fig. 2). The domains of *Hox1* and *Mist* expression filled the inner area, whereas the lateral expression domains of *Hox4* laterally expanded inward and merged with each other at the middle of the shell field (Fig. 2G–J). Therefore, based on these expression patterns, we identified the following three distinct cell populations in the larval shell field of *N. fuscoviridis*: population I, positive for *Hox1*, *Mist* and *Hox4*; population II, positive for *Hox1* and *Mist*, but negative for *Hox4*; and population III, positive for *CS1* and *dpp*. *Tyr* expression was detected in the area of population II.

### Characterization of the shell field cells examined from the single-cell transcriptome analysis

To further characterize the shell field cell populations, we performed a single-cell transcriptome analysis of 8 hpf larva. We first conducted bulk transcriptome analysis of multiple developmental stages and adult organs and constructed a reference gene model (see Methods for details) with high quality and low redundancy, as BUSCO completeness was 94.4% (single: 88.7%, duplicate: 5.7%, fragmented: 1.6%, missing: 4.0%). We used this transcriptome-based gene model as a reference for subsequent single-cell transcriptome analysis, as was performed in other non-model animals (34). We constructed a single-cell transcriptome library based on the 10X chromium system and sequenced the library using the HiSeq X Ten platform. Subsequent single-cell transcriptome analysis using the CellRanger (26) and Seurat packages (28) in R identified 14 cell clusters (Fig. 3A; marker genes for each cluster are shown in Fig. S2–3 and Table S2; see Methods for details). Among these clusters, clusters 4, 7, and 8 showed the gene expression profiles of the shell field genes examined in this study (Fig. 3B). *Hox1* and *Mist* were predominantly expressed in clusters 4 and 8, whereas *Hox4* was highly expressed only in cluster 4 (Fig. 3B). These expression profiles are consistent with those described above, and we annotated that population I corresponds to cluster 4 (*Hox1*, *Mist*, and *Hox4* are all positive), and population II corresponds to cluster 8 (positive for *Hox1* and *Mist* and negative for *Hox4*). *Tyr*-positive cells were found in cluster 8 (population II; Fig. 3B) but not in cluster 4. We also examined the expression of transcription factors that are known to be expressed in molluskan shell field cells (15, 20, 35–37) (Fig. S3). We found that engrailed gene and *Hox2* were predominantly expressed in cluster 8, whereas *Hox5* showed higher expression in cluster 4, and *Lox5* was expressed in both clusters. In addition, we detected and annotated the marker genes of each cluster and found other transcription factors in the marker genes for clusters 4 and 8 (Table S2); TALE homeobox genes *Pbx* and *Meis* were found as marker genes of cluster 4, and *Lhx3/4, SoxD and Ese* as marker genes of cluster 8 (Fig. S2).

**Figure 3.**
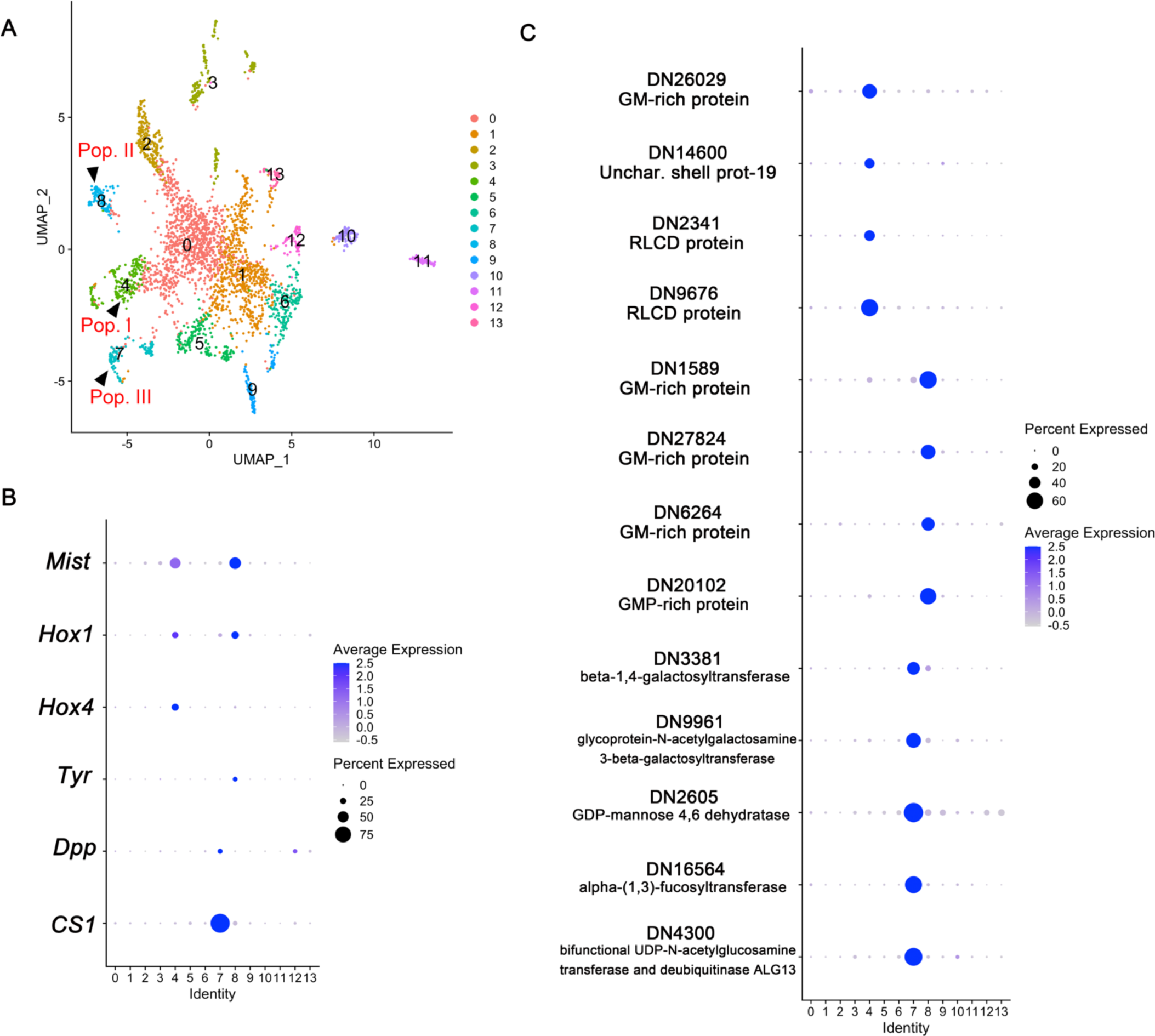
Single-cell transcriptome analysis of 8hpf *Nipponacmea fuscovilidius*. (A) The UMAP plot showing the cell clusters of 8hpf limpet. (B-C) Dotplot showing the expression of shell filed genes (B) and selected marker genes for cluster 4, 7 and 8 (C).

Clusters 4 and 8 were characterized by the expression of glycine- and methionine-rich proteins and repetitive low-complexity domain (RLCD) genes (Fig. 3C; Fig. S5, Table. S2). Hence, they were likely to be involved in the secretion of shell matrix proteins. Notably, there were differences in the repertoire of RLCD genes expressed in each cluster (Fig. 3C; Table S2), suggesting that functional differentiation occurred in populations I and II in the larval shell field. Population III was characterized by the expression of *CS1* and *Dpp* and was thus likely to correspond to cluster 7 (Fig. 3B). Several other known transcription factors expressed in the shell field, including *Lox5*, *GATA1*/*2*/*3*, *Pax2*/*5*/*8*, *Gsc*, and *Post2* were predominantly expressed in cluster 7 (Fig. S2, Table S2). In addition, the *AP2* and *Uncx* transcription factors were identified as marker genes for cluster 7 (Fig. S2; Table S2). Besides, this cluster is characterized by predominant expression of multiple types of enzymes for biosynthesis or modification of glycans and glycoproteins, such as glycoprotein-N-acetylgalactosamine 3-beta-galactosyltransferase, _α_-(1,3)-fucosyltransferase and, GDP-mannose 4,6 dehydratase (Fig. 3C). Together with the expression of chitin synthase, these cell populations may play other roles in shell development. Further characterization of the cell populations is currently underway in the authors lab, referring the single-cell transcript analyses of late-stage larvae and the adult mantle.

### The shell field cell differentiation in the isolated 2q blastomeres

Next, we investigated whether inductive signals from blastomere lineages other than 2q were required for shell field cell differentiation. We first isolated 1Q blastomeres from eight-cell embryos. After one round of division, 2q blastomeres were isolated and cultured individually until 9 hpf (Fig. S1). We found that signals for all six shell filed genes (*Hox1*, *Mist*, *Hox4*, *Dpp*, *CS1*, and *Tyr*) appeared in most isolated 2q fragments fixed at 9 hpf (Fig. 4A–F). Although we could not distinguish the characterization of 2q blastomeres along the dorsoventral axis (namely, 2a, 2b, 2c, or 2d) at the 8–16-cell stage, it is notable that all shell filed genes examined showed positive signals in most of the isolated fragments (Fig. 4). The results indicated that 2q blastomeres show an equivalent potential to activate shell field genes at the 16-cell stage. The expression of those marker genes, however, were not detected in any cell clusters that developed from the isolated 2Q macromeres (Fig. 4G–L). Therefore, cell-cell interactions between 2q descendant cells and other lineages, such as 2Q and its descendants, are not required for the expression of the above shell field genes.

**Figure 4.**
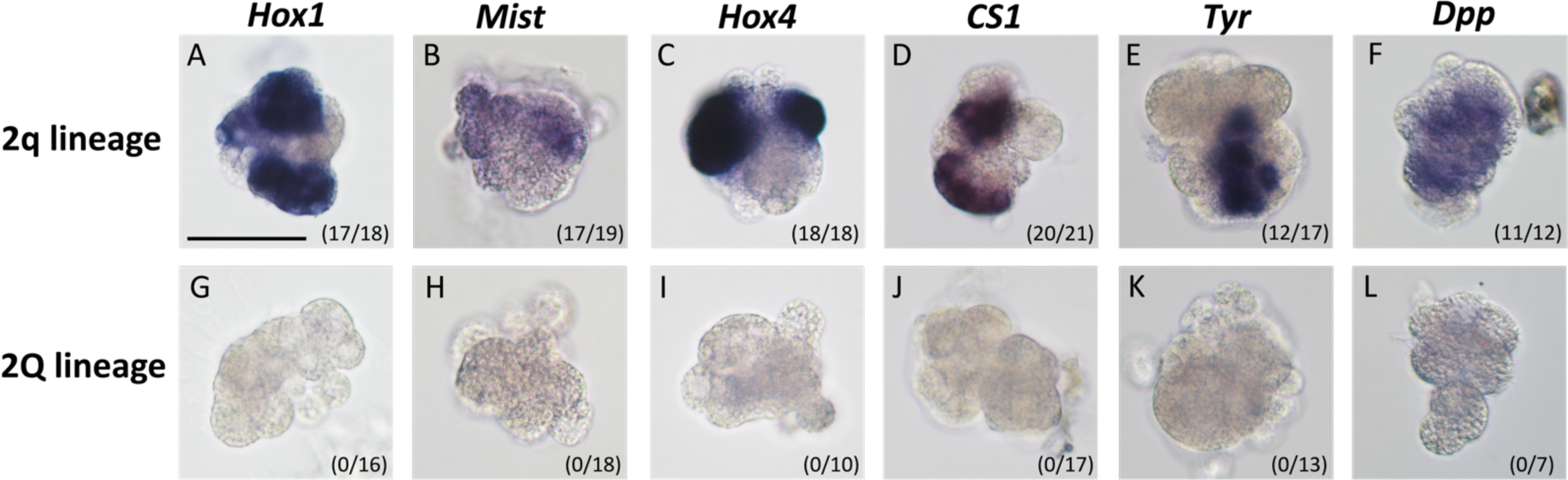
Shell field cell differentiation in the isolated 2q. Expression patterns of shell field-genes in embryos derived from 2q and 2Q isolates. Scale bar = 50 μm. Numbers indicate the number of embryos in which expression was detected (left) or observed (right).

Notably, no shell field genes were ubiquitously expressed in the isolated fragments, suggesting that the spatial expression of these genes may be locally regulated among ectodermal cells within the 2q lineage. Nevertheless, in most experiments so far, we found that these isolated cell clusters barely survived long after 9 hpf, possibly due to the damage caused by experimental manipulation. Therefore, examining other morphogenetic events, such as cell movements or shell matrix secretion, in fragmented embryos remains challenging.

### Shell field gene expression in cytokinesis-arrested embryos

Next, we determined how the 2q descendant cells acquired specific gene expression in each population (I–III) without interacting with other quartet lineages. As the blastomeres isolated after the 16-cell stage were too fragile to proceed with further development, we used an alternative method: blocking cytokinesis with the actin polymerization inhibitor, cytochalasin-B. These cytochalasin-B-treated embryos underwent nuclear proliferation without further cytoplasmic division and were fixed at 9 hpf (Fig. S6).

To determine whether shell field cell differentiation still occurred after cytokinesis arrest at the 16-cell stage, the expression of six shell field genes was assessed. In the cases of *Hox1* and *Mist*, signals for both genes were detected in the four syncytial blastomeres of the cytokinesis-arrested embryos examined (20/20, 18/18; Fig. 5A–B, G–H). Considering their size and position (Fig. S6), we determined that the *Hox1*- and Mist-positive syncytial blastomeres were 2q descendants. This is consistent with the above results that *Hox1* and *Mist* is positive in the isolated 2q descendants, but not in the isolated 2Q descendants. In contrast, when *Hox4*, *CS1*, and *Dpp* were used as markers, only approximately half of the embryos showed a positive expression signal (11/22, 33/63. 18/28; Fig. 5C–F, I–L). Notably, even in positive embryos, none showed expression in all four syncytial blastomeres (0/22, 0/63, and 0/28, respectively; Fig. 5). This is in contrast to *Hox1* and *Mist* which were positive in all four syncytial cells in most of the observed embryos (18/20 and 18/18, respectively). None of the treated embryos showed *Tyr* expression (Fig. 5E). These results are consistent with the expression profiles of shell field genes; the onset of shell field expression was slightly later for *Hox4*, *dpp* and *CS1* compared with that of *Hox1* and *Mist*, and *Tyr* expression commenced at the latest stage. These results indicate that cytokinesis after the 16-cell stage is not required for the expression of *Hox1* and *Mist*. On the other hand, it is suggested that additional cytokinesis dependent regulatory inputs are required for *Hox4*, *Tyr*, *dpp* and *CS1* expression.

**Figure 5.**
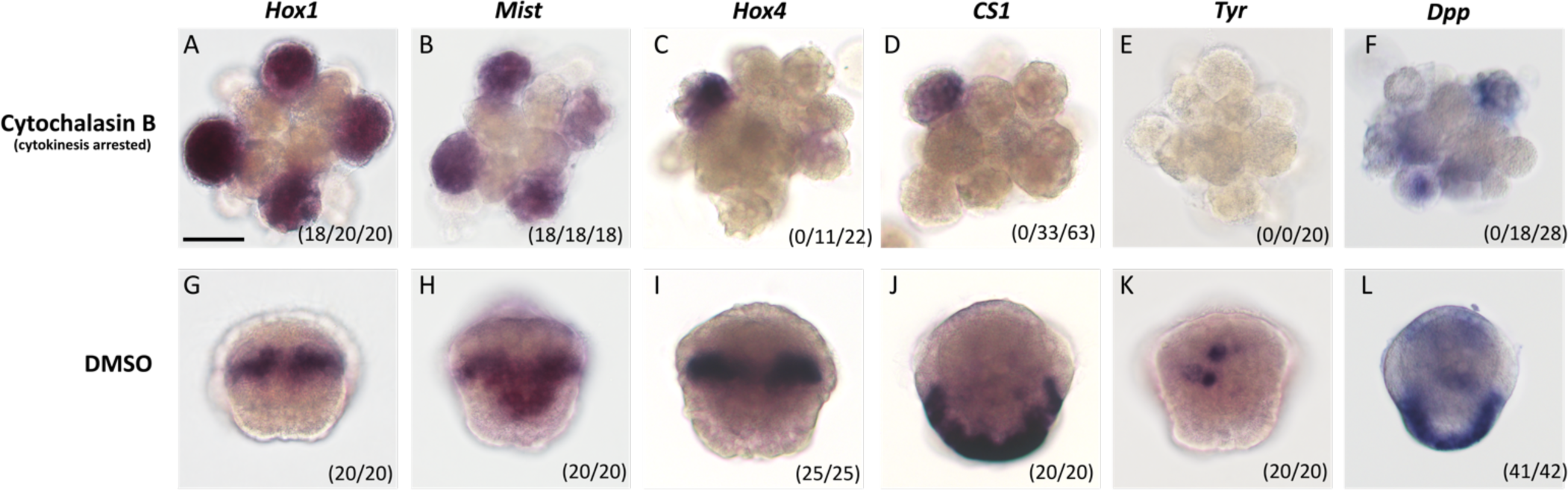
Shell field genes expression in cytokinesis-arrested embryos. Cytokinesis-arrested samples are shown in the vegetal view, whereas the controls are shown in the dorsal view. The numbers in A–F indicate the number of embryos in which expression was detected in the four syncytia (left), expression was detected at least one syncytia (middle), and observed (right). Numbers in G–L indicate the number of embryos in which expression was detected (left) or observed (right).

## Discussion

### Three subpopulations of the limpet larval shell field

Based on the spatiotemporal expression of the six shell field genes, the developing shell field appeared to be composed of at least three cell populations that formed at different points of development and displayed distinct cell behaviors during shell field morphogenesis. The shell field was first visible by the expression of *Hox1* and *Mist* on both the dorsal and ventral sides at 5 hpf (Fig. 1A, A’, G, G’). Subsequently, expression was gradually restricted toward the dorsal side (Fig. 1B–E, B’–E’, H–L, H’–L’). Although such expression dynamics may simply suggest a shift of gene expression to the dorsally localized cells, Yang et al (33) revealed that this change in signals was, in fact, underlain by the lateral cell movement toward the dorsal area. This is consistent with cell lineage studies of gastropods showing that the shell field is derived not only from the dorsal ectodermal cells of 2d, but also from the ventral and lateral cells (2a, 2b, 2c).

After 7 hpf, three distinct cell populations were observed in the dorsal areas (Figs. 1 and 2). *Hox4* expression is detected in the subpopulation of the *Hox1*-positive area; thus, the two populations are now recognized as population I (positive for *Hox1*, *Mist* and *Hox4*) and population II (positive for *Hox1*-*Mist* and negative for *Hox4*). The remaining population was characterized by *dpp* and *CS1* expression (cell population III). These *CS1*-expressing cells were first found immediately next to the bottom border of cell population II. In later stages, they gradually encircled most of the shell field region from the bottom and became the outer edge of the shell field (Fig. 1 Y–CC, Fig. 2).

We identified several effector genes for each shell field population using single-cell transcriptome analysis. Populations I and II expressed genes encoding glycine- and methionine-rich proteins and RLCD-containing proteins, whereas population III was characterized by the expression of chitin synthase and multiple types of enzymes for modification or biosynthesis of glycan and glycoprotein (Fig. 3; Table S2). The expression of distinct effector genes suggests that each population plays a distinct role in the development of the shell field, although further characterization of each population requires additional studies.

In other mollusk species, different subgroups of developing shell field cells have been reported (12, 18, 36, 38–40). In *L. stagnalis*, Timmermans (38) uncovered the distinction between the peripheral and central groups of shell field cells that became distinguishable after shell field invagination based on the activities of alkaline phosphatase, Tyrosinase, and peroxidase. As the mantle tissue of adult animals was previously subdivided into different zones that perform specific functions in the shell-forming process, Timmermans also pointed out that this mantle zonation was likely to be established as early as the shell field first appeared in an embryo. Herlitze et al. (18) supported this idea by showing the similarity in the gene expression profiles between the central region of the larval shell field and the inner zones of the adult mantle, as well as the similarity between the outer region of the larval shell field and the outer zones of the adult mantle. However, because we found little overlap between the gene repertoires of their study and ours, to complete our knowledge regarding the embryonic shell field cells and mantle zonation, systematically characterizing all shell field cell populations and determining homologous cell groups across mollusk species remains crucial.

### Autonomous development of the shell field in 2q cells

Although 2q is known to be the cell lineage that gives rise to the entire shell field in mollusks, whether these 2q cells already possess the potential to differentiate into the shell field on their own is debatable. Although Raven (10) originally proposed that the acquisition of the shell field fate did not occur until the tip of the archenteron made contact with the ectoderm during gastrulation, other authors (13, 41) highlighted that interactions with any endodermal cells, or even the mesentoblast or the polar lobe, were sufficient to induce the ectoderm to become the shell field.

Nevertheless, the lines of evidence revealed in the current study are not consistent with previous views on shell field induction. First, and most importantly, when the 2q cells were isolated from 2Q macromeres and cultured separately, the independently developed 2q fragments continued to differentiate and activate the expression of six shell field-related genes (Fig. 4). Given that it was technically feasible to rear the isolated fragments on the stage-shell matrix, our study did not exclude the possibility that there might still be some other developmental events of the shell field cells depending on their interactions with the endoderm or other cell lineages. Indeed, several previous studies showed that the disruption of the interactions between the shell field-forming 2q and other blastomere lineages commonly resulted in embryos with abnormal shells (16, 42–44). For example, Labordus & Van der Wal (42) demonstrated that the early ablation of the 3D macromere during the first 60 min after the fifth cleavage frequently caused the failure of external shell formation and an increase in the underdeveloped internal shell masses in *T. obsolete*. Moreover, McCain (16) revealed that the interactions within and between the second and third quartets were required to develop normal external shells in the same gastropod species. However, our study indicated that the initial process of shell field establishment does not require any interactions between 2q blastomeres and other lineages to activate the expression of shell field-genes.

According to Raven’s induction model, specified shell field cells appear specifically on the dorsal side, where the tip of the archenteron contacts the overlying ectoderm. Therefore, the shell field gene expressions should initially be detected at a single spot on the dorsal posttrochal area before they later expand in all directions as the shell field develops (12, 20). In contrast to this induction scenario, the early expressed shell filed genes(i.e., *Hox1* and *Mist*) indicated that the shell field cells did not specifically emerge from the dorsal cells, but rather from the cells on both the lateral and ventral sides (Fig. 1A, A’, G, G’). During the later stages, these shell field cells became localized to the dorsal area where the shell field developed; therefore, the specification event was likely to occur before dorsal migration (Fig. 1D–F’, J–L’). This notion is supported by the results of cell movement inhibition where the trochophore larvae of the limpet *L. goshimai* were treated with blebbistatin, which prevents the phosphorylation of non-muscle myosin II, and the dorsal migration and other morphogenic events of shell field formation were inhibited (45). However, the differentiation of the shell field cells was not interrupted, as indicated by the signals of the shell field genes *BMP2/4* (*Dpp*), *GATA2/3*, *Hox1* and engrailed. The signals of these genes expanded on both lateral sides in the blebbistatin-treated group, whereas the signals in the control group were restricted to the dorsal area. Moreover, unlike *L. stagnalis*, we noticed that the limpet *N. fuscoviridis* does not form an invaginated archenteron that forms a single contact point with the dorsal ectoderm. Thus, most ectodermal cells were in direct contact with the underlying endoderm, as previously shown by Hashimoto et al. (15). Again, this observation agrees with the argument of several previous authors that the formation of a deep archenteron is probably not a common feature of mollusk development (11, 14, 33). Overall, these findings are not consistent with the inductive roles of other cell lineages (i.e., 1q^1^, 1q^2^ and 2Q) in initiating shell field cell specification after the 16-cell stage.

### Differentiation of the shell field populations

Six shell field genes were expressed in the isolated 2q fragments. In contrast, a clear difference in expression was observed between the two groups of shell field genes in cytokinesis arrest experiments. In the cases of *Hox1* and *Mist*, the expression was observed in both isolated 2q and cytokinesis-arrested embryos (Figs. 4 and 5). Thus, the expression of *Hox1* and *Mist* is regulated in a more cell-autonomous manner in the 2q lineages. However, some cells derived from the isolated 2q were negative for *Hox1* or *Mist*. Therefore, although the expression of *Hox1* and *Mist* does not require cytokinesis after the 16-cell stage, additional mechanisms must exist to restrict *Hox1* and *Mist* expression in some 2q derived cells.

In contrast to *Hox1* and *Mist*, our results suggest that the expression of *Hox4*, *dpp*, *CS1,* and *Tyr* more dependent on cytokinesis after the 16-cell stage; fewer embryos were positive for the 16-cell cytokinesis-arrested embryos (Figs. 4 and 5), although all of them showed expression in the isolated 2q fragments. These results suggest that although initial cell differentiation toward the precursors of populations I and II (positive for *Hox1* and *Mist*) occurs in a more autonomous manner in 2q derivatives, further differentiation into populations I and II, as well as the differentiation of population III, requires additional mechanisms. One plausible explanation for this observation is that cytokinesis arrest inhibits cell-cell interactions among the 2q derivatives. Although, cell-cell interactions among 2q-derived populations in 2q-isolated fragments occur normally, cell-cell interactions may not occur normally in the syncytia. The expression of *Hox4*, *dpp* and *CS1* was not completely suppressed in cytokinesis-arrested embryos, possibly because the signal from one of the 2q syncytia reached the other 2q syncytium. Although this is not the sole possible explanation for the observed results, we can safely conclude that some of the shell field-related genes (*Hox4*, *dpp*, *CS1*, and *Tyr*) show clear differences in their dependence on cytokinesis after the 16-cell stage compared with that of *Hox1* and *Mist*. Therefore, we concluded that shell field cell differentiation into functional populations occurs in multiple steps. In this context, it is notable that *dpp* was reported to be required for the activation of other marker genes in populations III such as *CS1*, although the expression of *Hox1* and engrailed genes was not affected (15). Thus, Dpp signaling is likely involved in further differentiation of population III.

Our results on 2q fragments and cytokinesis-arrested embryos provide the first step toward understanding the entire process of shell field development. Elucidation of the shell field differentiation mechanisms will open the window toward a comprehensive understanding of the evolutionary innovation of shell plates. In addition, it will allow us to address the question of the evolutionary link between shell plates and spines of aplacophora and chitons.

## Acknowledgements

The authors would like to thank Dr. Satoshi Yamazaki and Rei Hirochika for kind support for using 10X Chromium controller. The authors would like to thank Maitreyee Pargaien and Li Yuanhong for gene isolation. This work was supported by the Japan Society for the Promotion of Science KAKENHI grants 18H04812 (Grant-in-Aid for Scientific Research on Innovative Areas), 18K14762 (Grant-in-Aid for Young Scientists), and 23K05873 (Grant-in-Aid for Scientific Research (C)) to YM and 18H04004 (Grant-in-Aid for Scientific Research (A)) to HW.

## List of Supplementary Materials

**Text S1** Methods for bulk transcriptome analysis

**Text S2** Annotation of each cluster in single-cell analysis

**Fig. S1** Step-wise blastomere isolation protocol

**Fig. S2** Expression of the marker genes and transcription factor genes in single-cell transcriptome analysis

**Fig. S3** *In situ* hybridization of selected marker genes in each cluster

**Fig. S4** Molecular phylogeny of transcription factors

**Fig. S5** RLDC and GM-rich protein sequences in cluster 4 and 8 marker genes

**Fig. S6** The 9 hpf embryos blocked in cytokinesis from the 16-cell stage

**Table S1** Primers used for gene isolation

**Table S2 (separated file)** Marker genes and annotations for each cluster

**Dataset S1 (separated file)** Aligned sequences of transcription factor for phylogenetic analysis

