## Supplementary material for "Development of shell field populations in gastropods": Text S1-2, Fig.S1-6, Table S1

### List of Supplementary Materials

**Text S1** Methods for bulk transcriptome analysis

**Text S2** Annotation of each cluster in single-cell analysis

**Fig. S1** Step-wise blastomere isolation protocol

**Fig. S2** Expression of the marker genes and transcription factor genes in single-cell transcriptome analysis

**Fig. S3** *In situ* hybridization of selected marker genes in each cluster

**Fig. S4** Molecular phylogeny of transcription factors

**Fig. S5** RLDC and GM-rich protein sequences in cluster 4 and 8 marker genes

**Fig. S6** The 9 hpf embryos blocked in cytokinesis from the 16-cell stage

**Table S1** Primers used for gene isolation

**Table S2 (separated file)** Marker genes and annotations for each cluster

**Dataset S1 (separated file)** Aligned sequences of transcription factor for phylogenetic analysis

**Text S1 Methods for bulk transcriptome analysis**

Total RNA was extracted from egg, eight-cell stage (1 h and 35 minutes post fertilization [mpf]), 16-cell stage (1 h and 55 mpf), 32-cell stage (2 h and 20 mpf), morula (3 h and 10 mpf), gastrula (4 hours post fertilization [hpf]) swimming gastrula/ early trochophore larva (4 h and 50 mpf)), trochophore larva (10 hpf), veliger larva stages (17 hpf and 25 hpf), and adult organs (mantle, head including internal organs, foot, ovary including associated internal organs, and testis). RNA extraction from mantle, foot, ovary, 17hpf and 26hpf larvae was performed using NucleoSpin ® RNA Plus XS (TAKARA), while extraction from other tissues/embryos was performed using Trizol and Purelink Dnase Set (Invitrogen). For eggs - 10 hpf samples, RNA was extracted from two batches. Library construction and sequencing on NovaSeq 600 was performed by Novogene for Egg-10 hpf samples and by Macrogen for veliger larvae and adult organ samples. Raw sequences were deposited in the DNA Data Bank of the Japan Sequence Read Archive (PRJDB18060).

**Text S2 Annotation of each cluster in single-cell analysis**

We annotated the cell population of each cluster in the single-cell RNA sequencing (RNA-seq) analysis. This was based on annotations of the detected marker genes for each cluster (Table S2) and observations of their expression patterns by *in situ* hybridization, as described below.

**Cluster 2: Ciliary cells**

Cluster 2 was annotated as the ciliary cells. Many of the marker genes were genes known to be generally involved in ciliogenesis, such as α-tubulin, β-tubulin, and tektin (Table S2). Indeed, one of the marker genes, tektin, was expressed in the ciliated cells of trochophore larvae (Fig. S2–3). Therefore, we concluded that cluster 2 corresponded to ciliated cells. Considering that the apical tuft cells are presumed to belong to cluster 3 or 12 (as discussed below), cluster 2 corresponds to the ciliary cells constituting the prototroch.

**Cluster 12: Anterior ectoderm**

Cluster 12 was annotated as the pretrochal anterior ectoderm. One of the marker genes, *FoxQ2c*, is expressed in the apical region at 8 hpf (1) (Fig. S3). *dpp* (also highly expressed in the shell field) and *Dlx* (also highly expressed in the posterior ectoderm) were expressed in cluster 12 in single-cell transcriptome analysis (Fig. 3 and Fig. S2–3). We conducted *in situ* hybridization of these genes and revealed that they were expressed in the nonapical anterior ectoderm (Fig. 1 and Fig. S3). Therefore, we judged that cluster 12 corresponds to the pretrochal anterior ectoderm.

**Cluster 3: Neuronal cells**

Cluster 3 was annotated as the neuronal cells. Marker genes included several calcium channels and synaptotagmin, which are generally involved in neuronal functions. Indeed, some marker genes (leucine-rich repeat-containing protein 74 [*LR74*] and uncharacterized protein [*Unchar-2260*]) were expressed in the ectoderm near the apical region (Fig. S3). *LR74* was also expressed in a few cells in the posterior ectoderm (Fig. S3). Both regions correspond to the position of neuronal cells (anlage of the cerebral ganglion and posterior larval sensory organ) in the trochophore of mollusks (2). Therefore, we considered cluster 3 to be neuronal cells.

**Cluster 6 Posterior ectoderm**

We performed *in situ* hybridization of transcription factors *Sp6-9* and *Dlx*, which are highly expressed in cluster 6 in single cell transcriptome analysis (Fig. S2). We found that those gene were expressed in the posterior ectoderm (Fig. S3). Therefore, cluster 6 was annotated as the posterior ectoderm.

**Cluster 9 Ventral ectoderm**

We examined the expression pattern of *FoxAB*, a marker transcription factor for cluster 9 (Fig. S2), and revealed that *FoxAB* specifically expressed in the posttrochal ventral ectoderm at 8 hpf larvae (Fig. S3). Thus, Cluster 9 was identified as the posttrochal ventral ectoderm.

**Cluster 11: Cells around mouth**

Cluster11 was considered to be the cells around the mouth because *Erg*, the marker transcription factor of cluster 11 (Fig. S2), showed expression in cells around the mouth by *in situ* hybridization (Fig. S3).

**Cluster 5: Mesoderm**

Cluster 5 was annotated as the mesoderm. Marker genes included transcription factors *Twist,* which are known to be expressed in the mesoderm of the limpet trochophore (Table S2; (3)). Additionally, the marker gene *Alx* was expressed in the anterior mesoderm at 8 hpf larvae (Fig. S3). Thus, cluster 5 was identified as the mesoderm. It should be noted that it is unclear whether posterior mesoderm cells are included in this cluster or in other clusters.

**Cluster 10 and 13: Endoderm**

Marker genes in cluster 13 included *Cdx*, a transcription factor known to be expressed in the hindgut of a wide range of animals, including mollusks (4, 5) (Table S2). Moreover, one of the marker transcription factor genes, *Blimp1*, was expressed in the endoderm in limpet (Fig. S3). We also examined the expression of two marker genes of cluster 10 (tryptophan 2,3-dioxygenase (*T23O)* and kynurenine 3-monooxygenase (*KMO)*) and found that both genes were expressed in the endoderm (Fig. S3). Therefore, we concluded that these two clusters were endodermal. Differences in the distribution and properties of the cells constituting these two clusters are currently unknown.

**Cluster 0 and 1: Unknown**

Only a few marker genes were detected in clusters 0 and 1 (Table S2). Furthermore, they were also expressed in other clusters (Fig. S2B and Table S2), most of which were housekeeping or unannotated genes. (Table S2). Therefore, the names of clusters 0 and 1 were set as “Unknown.”

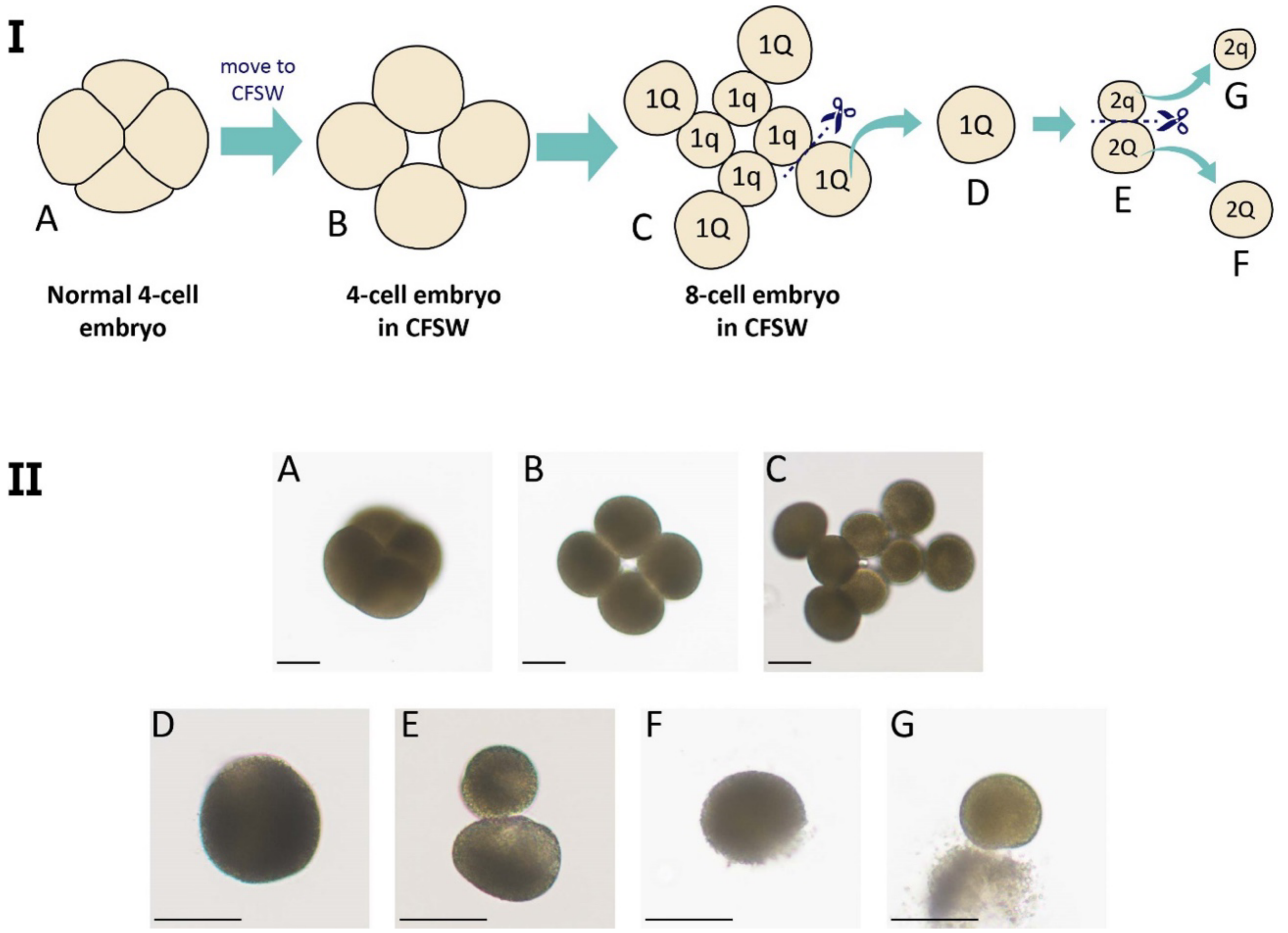


**Fig. S1 Step-wise blastomere isolation protocol**

The procedure for blastomere isolation is displayed in the summary diagram above (I), whereas pictures of the actual embryos or cells are shown below (II). The four-cell embryo in ASW (IA, IIA), the four-cell embryo after treatment with the dissociating solution and being moved to CFSW (IB, IIB), the 8-cell embryo in CFSW (IC, IIC), the isolated 1Q cell (ID, IID), the two daughter cells of the 1Q cell (IE, IIE), the isolated 2Q cell (IF, IIF), and the isolated 2q cell (IG, IIG). Scale bar = 50 µm. Abbreviations: ASW, artificial seawater; CFSW, calcium-free seawater.

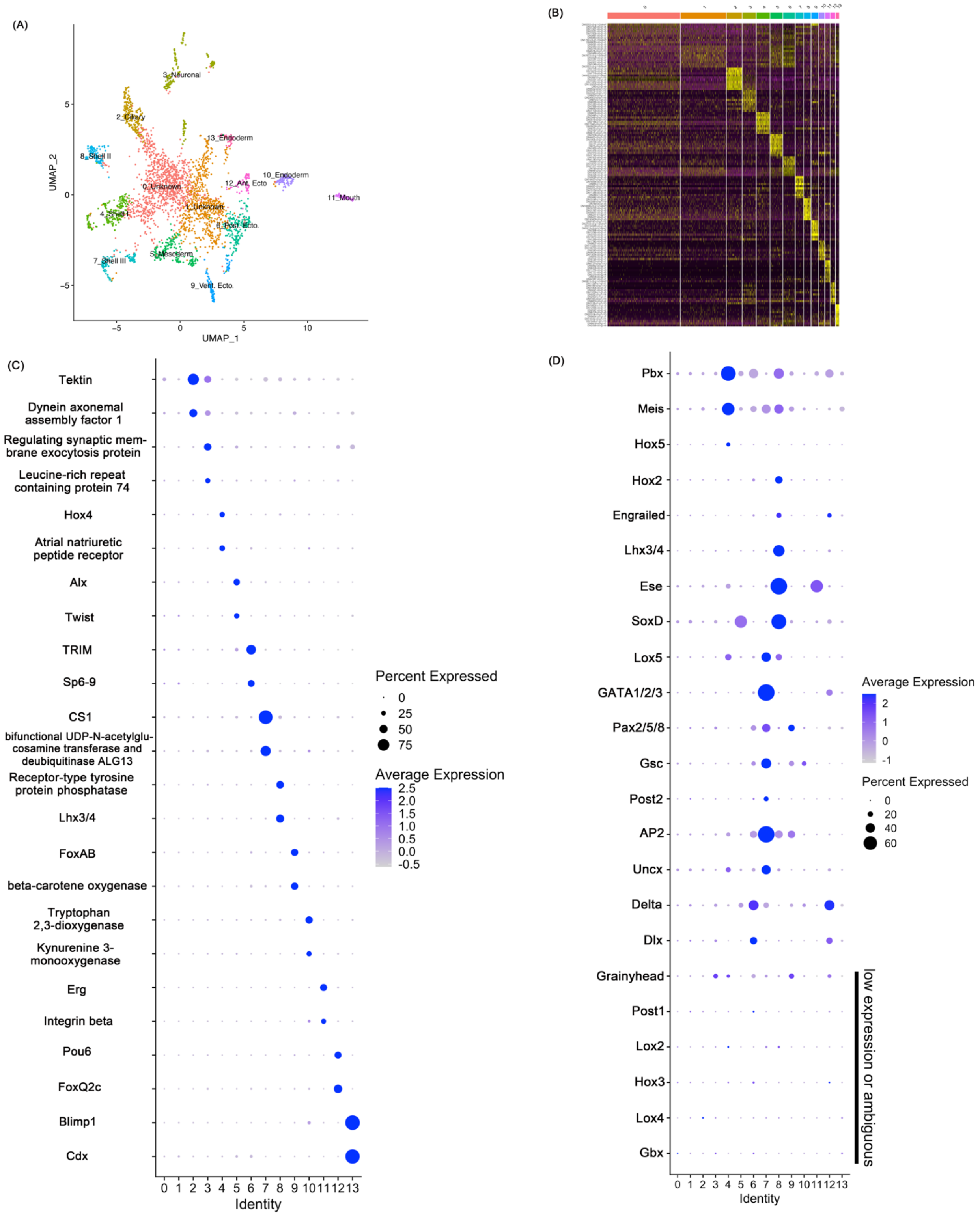


**Fig. S2 Expression of the marker genes and transcription factor genes in single-cell transcriptome analysis**

**Fig. S2 Expression of the marker genes and transcription factor genes in single-cell transcriptome analysis**

**(A)** The UMAP plot showing the cell clusters of 8 hpf limpet with annotation. (B) Heatmap of marker gene expression in each cluster. (C) Dotplot of selected marker genes in each cluster. (D) Dotplot of transcription factors which are known to be expressed in shell field or marker genes for cluster 4, 7 or 8.

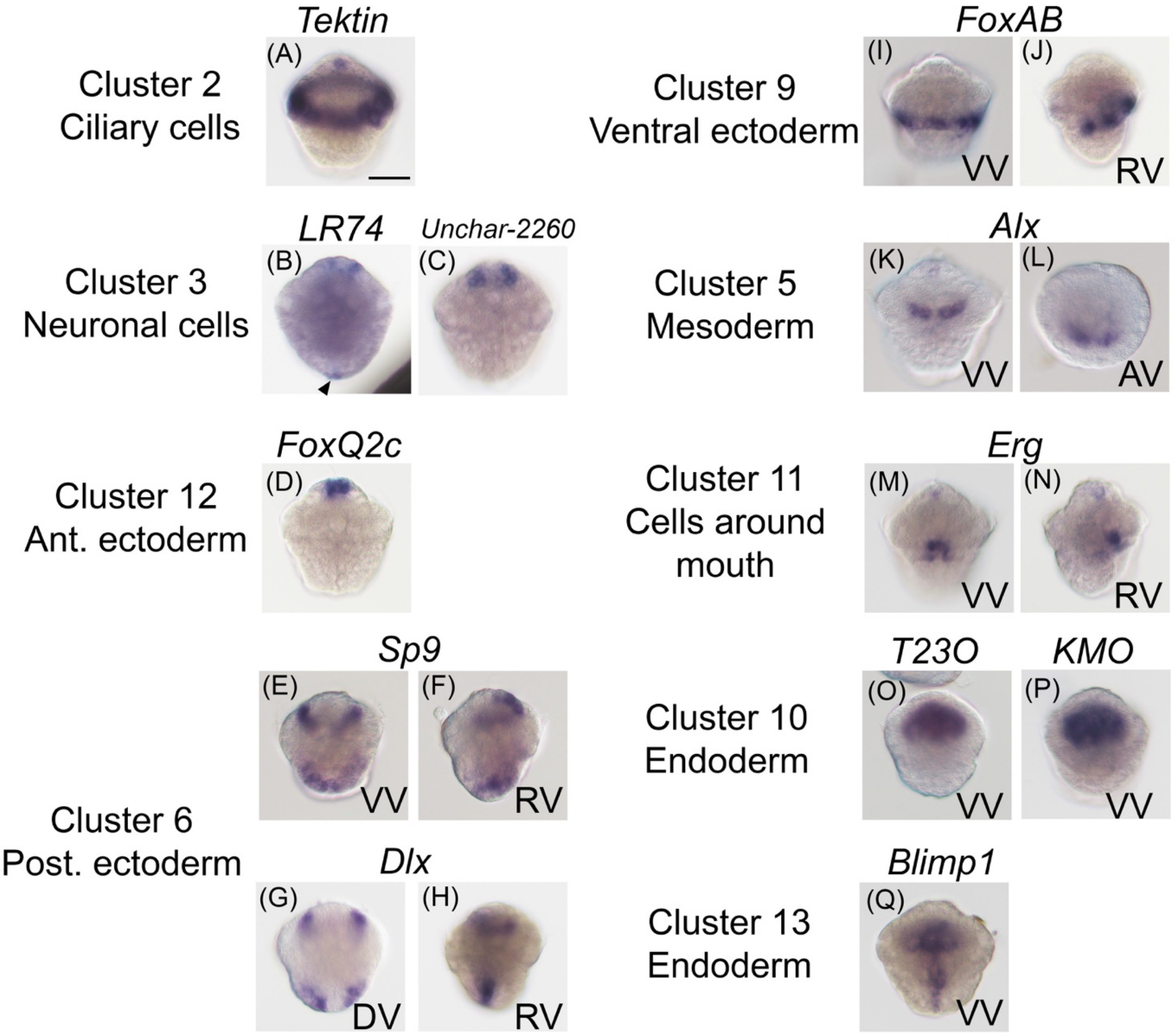


**Fig. S3 *In situ* hybridization of selected marker genes in each cluster**

The expression patterns of selected marker genes in each cluster are shown. (A) *Tektin* was expressed in the prototroch and apical cells. (B, C) Expression of *LR74* and *Unchar-2260* was detected in ectodermal cells near the apical region. *LR74* expression was also detected in a few cells of posterior ectoderm. (D) *FoxQ2c* expression in the apical ectoderm. (E, F) *Sp6-9* is expressed in the part of the posterior ectoderm. Note that even the high expression was not detected except for cluster 6 in the single-cell transcriptome analysis, expression in the pretrochal ectoderm was detected. (G–H) *Dlx* expression was detected in the dorsal part of the posterior ectoderm and pretrochal ectoderm. (I-J) *FoxB* was expressed in the posttrochal ventral ectoderm. (K–L) *Alx* is expressed in the anterior mesoderm and apical cells. (M–N) *Erg* is expressed in the cells around mouth. Subtle expression in apical cells was also detected. (O-P) Both *T23O* and *KMO* were expressed in endodermal cells. (Q) *Blimp1* expression was detected in endodermal cells. VV, ventral view; DV, dorsal view; AV, animal view; RV, right side view. Scale bar = 50 μm.

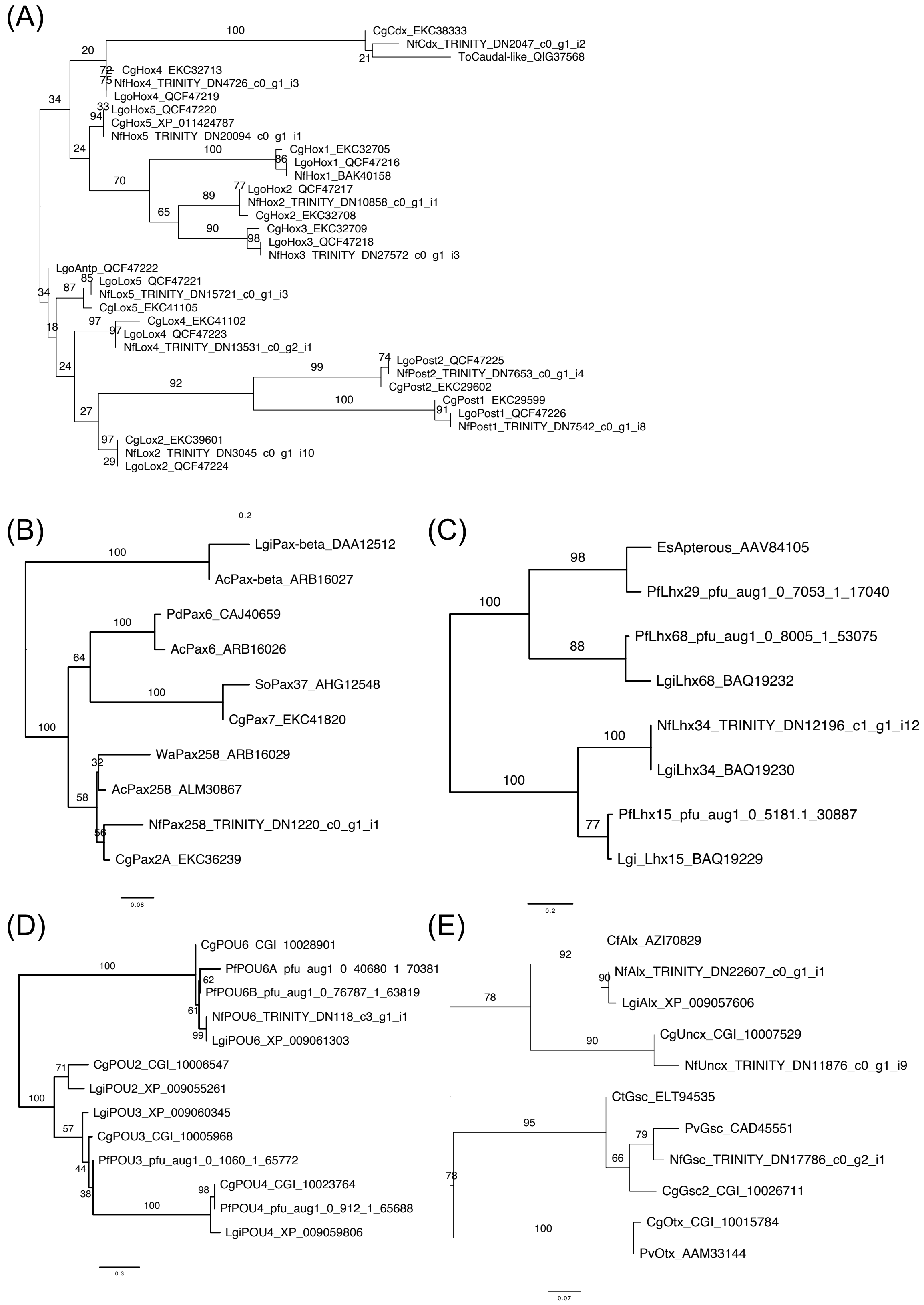


**Fig. S4-1 Molecular phylogeny of transcription factors**

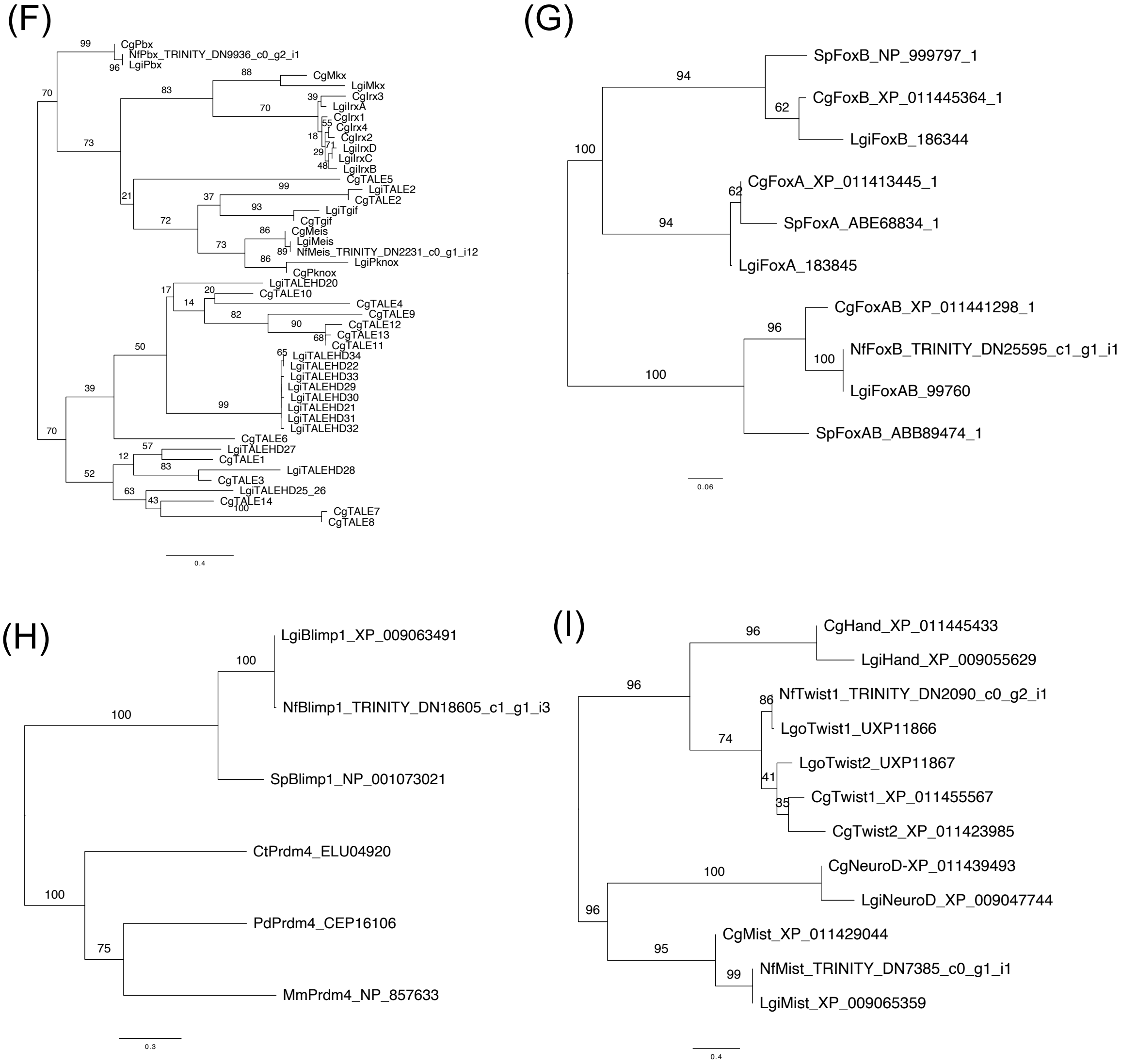


**Fig. S4-2 Molecular phylogeny of transcription factors**

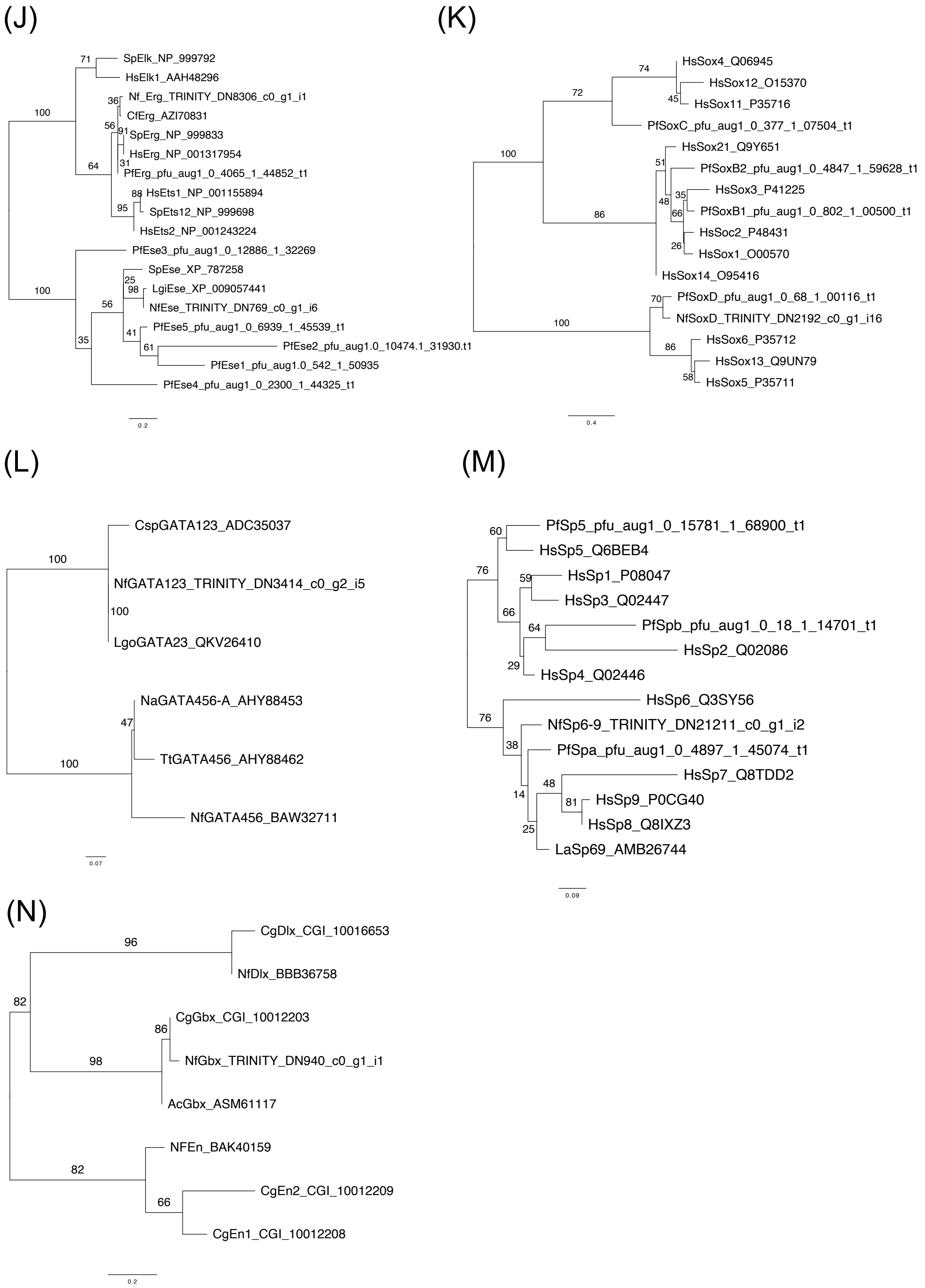


**Fig. S4-3 Molecular phylogeny of transcription factors**

**Fig. S4 Molecular phylogeny of transcription factors**

Maximum likelihood trees of transcription factors for orthology confirmation. Numbers in the nodes indicate bootstrap values of 1000 replicates. The trees were visualized by FigTree (http://tree.bio.ed.ac.uk/software/figtree/). (A) Hox and Parahox genes. LG + G model was used. (B) Pax genes. The LG + G4 model was selected. The annotations of spiralian Pax genes are depends on Wollesen et al. (6). (C) Lim homeobox genes. The LG + G4 model was used. The annotation of the molluscan Lim homeobox gene referred to Morino et al. (7). (D) POU homeobox genes. The LG + G4 model was selected. Annotation of molluscan POU homeobox genes followed Morino et al. (7). (E) PRD homeobox genes. The LG + G4 model was used. The annotation of the molluscan PRD homeobox gene referred to Paps et al (8). (F) TALE homeobox genes. The LG + G4 model was used. The annotation and sequence of the molluscan TALE gene were derived from previous research (9). (G) Fox genes. The LG + G4 model was selected. The annotations of molluskan Fox genes are depends on Yang et al. (10). (H) PRDM genes. DCMUT+G4 model was used. (I) bHLH genes. The LG + G4 model was used. The annotation of the molluscan bHLH gene referred to Bao et al.(11). (J) Ets genes. The LG + G4 model was selected. Annotation of molluscan Ets genes followed Koga et al. (12). (K) Sox genes. The LG + G4 model was used. The annotation of the molluscan Sox gene referred to Koga et al. (12). (L) GATA genes. The FLU+I model was selected. (M) Sp gene. The DAYHOFF+G4 model was used. The annotations of Sp genes are depends on Koga et al. (12). (N) Antp class homeobox genes. The LG + G4 model was used. Annotation of oyster homeobox genes followed Paps et al. (8). Cg, *Crassostrea gigas*; Nf, *Nipponacmea fuscoviridis*; Ct, *Capitella teleta*; Lgi, *Lottia gigantea*; Lgo: *Lottia goshimai*; Pd: *Platynereis dumerilii*; Tt: *Terebratalia transversa*; Na: *Novocrania anomala*; Csp: *Chaetopterus sp. MB-2010a*: Ac: *Acanthochitona crinite*; So: *Sepia officinalis*; Wa: *Wirenia argentea*; Sp: S*trongylocentrotus purpuratus*; Es: *Euprymna scolopes*; La: *Leptochiton asellus*; Cf: *Crepidula fornicata*; Pv, *Patella vulgate*; Mm: *Mus musculus*; Hs: *Homo sapiens*.

### Cluster 4 marker genes

TRINITY_DN14600_c0_g1_i1_Uncharacterized shell protein-19.prot (244 aa)

MLSLWAIGLLGLLNQVGAQQWGAAQSFQYTPPTQQTPTPANAQFGQMNKFFQRPIQTPSWGGASP
1 10 20 30 40 50 60

NPTPQNAWNFRQPNNMAPNVPNVPGAMPNMPGNMPNTMMNNMRNMMGNMRPGVTPNIPGSRGFGS
70 80 90 100 110 120 130

GYPNNMCYAPPNLPYLCTEKGENQHKFTLKEVMRMLGDKNIEKIPYCVPGIQRVNCANYDLAQYI
140 150 160 170 180 190

WQRGYCYCKRYHSNMNAFKKYEPLRCVLNNCGACSQEFFIGKSRIVDCD
200 210 220 230 240 244

TRINITY_DN9676_c0_g1_RLCD protein.prot (401 aa)

MIGVIFVTVCLGLRGVLGYPGGAPSSACVYLLPKHGYPHFEPQASISPFELVFSSATYSPGQEME
1 10 20 30 40 50 60

VAIVSEASPFNGWLLQVRRVDTDMVVGQFVSSPVGSRIVSCNGQNNSITHSNSFMFPNTTLRAKW
70 80 90 100 110 120 130

IAPQEDVGNLILIGTVLKDYSTFWGPLESNEIPFVPDAPSQDTPTTQKPTTSSTTTIIPSTSTSS
140 150 160 170 180 190

PTTNSPSTTTVSPTTATQATTTTIQPPITTTTTKTTPQPTFSNTTTPQPTTTSSTTPPPRITSST
200 210 220 230 240 250 260

TTPPRITSSTTTPPRITSSTTTPPRITSSTTTQPTTPSSTTQKVTTKLPSTTTTPSPTTSTKSPN
270 280 290 300 310 320

QPMNSTTQKPVPTSTKQAQNQSTETEPSTQENSSSETVPFAATKDDSKDTSGNGSGTVTSQNILL
330 340 350 360 370 380 390

LLSLWALCWCK
400 401

TRINITY_DN26029_c0_g1_i1_GM-rich protein.prot (233 aa)

MSNIRKLTPISKSNCIDEDRLYKSLQFGMKSSLFSLAVVQLCFQCVVTSFINLCGMCNFCSVFSV
1 10 20 30 40 50 60

IVYIHLCVVYDILFKSISAFLPGMTPGAAGGLGGPGGFPGGSGGFTGGSGGGSPMSMMLPLLMRG
70 80 90 100 110 120 130

GMDMRTMMMMMSMNRGQGGMDSMLPFLLMRNGGSENMLPMLMMMKNKGGAGGGMNPMVMAAMMGG
140 150 160 170 180 190

NDNMMQMLPFLMMNKQQSGPPANVALPAGAANPASGPI
200 210 220 230 233

**Fig. S5-1 RLDC and GM-rich protein sequences in cluster 4 and 8 marker genes**

TRINITY_DN2341_c0_g1_i7_RLCD protein.prot (877 aa)

```
MVRTKNLSSCACILVFLSNVILAIQQGIQQPSNAAIQPSTSTGYVQHPGYGSSPYHPHHMPLQYG
MQPSPYMMPMMDPDFLDHRMDMAKDMAKQRYAMNMYRHSYSPMGMYPRYPYQQPNQFGPGHHSAP
MNIVGGRQGASPVATHSTQKSKPNSGSKRLSRKKRFVPMMPIPFAPFGGFSPMLGVLRGAAVAGN
MAMVHRGRLANMRALAARRNRNGFNQNSRTRESSVSSFNSQPTRPMDNSFNAINNVVARSTGAQP
TVVQQLVPQPLSQPVMTQNIISQPVQRVELDPLLSQPALTQNLVLDQPSFIDPMGIGNQQQIIVD
NGLGGQQQAIVDGGLATQPQIMIDSGVVGQQPVVIDNTLAGQQQQIADQQAVNQQLAAQAASNKQ
LDAALQLQMDVQQAINQQTSQQSRLTDQQLTPQNTQQLYSDAGQQMDVLSGQQPQSSQQIYNLQQ
NQQALDQQAVNQQILNQQAFNQQALDQQTLNQQALIQQALNQPTLNQQVLDQQALNQQALNQQVL
NQQAFNQATLDQQTLNQQTLNQQAFNQQALDQQALNQQTLNQQAFNQPALGQQTFNQQTLNQPAL
NQQTLDQQAMNQQILNQQTLNQQAMDQQVLNQQILNGQSLNQQAFDQQAMNQQTMDQHAMKQQIL
NQQALNQQALDQQVLNQQILNGQALNQQAFNQPILNQQAFDISQSISGQQANSQQTTLTSNAPQS
VSNPYEGLNFQIQNVLGDLAASQTSSSQSNAVDVGVPQTLVQLQQPSNQQILVQQPTTQQGALQQ
STGQQMSLAYPDLFSFKDPLASNQGNSLQIGQAQAVEPATPGVVAAGGGGRPPGPPPPNPSQTGQ
QANLAALDLIANLASLNQRGVIPKHHGRPFNL
```

**Fig. S5-2 RLDC and GM-rich protein sequences in cluster 4 and 8 marker genes**

### Cluster 8 marker genes

TRINITY_DN1589_c0_g1_i12_GM-rich protein.prot (215 aa)

MNSLLSILSCCIGVATASFGMMNPMMGGMGSMLGMMGGFGGMGSMMGMHGFGGMGGMTGYPGITG
1 10 20 30 40 50 60

LSGLMGGFGGMNSMMGGMSGGMGGMDPMMGGGMGGGMGGMNPMMGGAMGGMNPMMGGAMGGMAGG
70 80 90 100 110 120 130

MSGMAGGMGGMSPMMGGVMGGMNPMMGGSMGGMNPMIGGAMGGMTGAMGGMNSMMGGGMGGMNPM
140 150 160 170 180 190

MGGGMGMMGGDMTGQNGFRK
200 210 215

TRINITY_DN6264_c0_g1_i9_GM-rich protein.prot (141 aa)

MYMNQYINTDSIDGKTITNRLYFKQSSSQPPTTKINMNTLFAIAVCFAFTATTLAFNPMMMGGMG
1 10 20 30 40 50 60

MMGGMGMMGGMGMMNPYMMGMGGLGMMGMGGMGMMGGMYNPMMMGYGGLGMLGGMGGYGMMGGML
70 80 90 100 110 120 130

GGMGGYGMMGK
140 141

TRINITY_DN20102_c0_g1_i18_ GMP-rich protein.prot (422 aa)

MPGPMGGMPGPMNGMHGPMGSMSGPMGSMPGPMGGMPGPMGGMPGSMGGMQGPVGGMPGQMSGSM
1 10 20 30 40 50 60

GGMQGPMGGMPGPMGGMPGSMGGMQGPVGGMPGQMSGPMGGMPMPADFQRPSGAWVDQSKLMAQQ
70 80 90 100 110 120 130

QRGDMGKIHDAQKKFMEKLIAIKTGTNGTANMTDEEKKQFLMSDGMPPMTPLQSAVGHEAEMVQN
140 150 160 170 180 190

MIEYEKQLAQAQHTVLKQFYTDVLTSRVVNTMFRKRLGEELQTLNALNKEVEKARKSMAGMTQQQ
200 210 220 230 240 250 260

LGVVPQPMMGGPQGIMMDPRASMIMSQSAMMGQMNPRAAMMGQMNPRAAMMSQMNPRAVMMGPMD
270 280 290 300 310 320

PRAAMMGQIDPRAAMMGQMDPRSAMMGQMNPRAAMMGQMDPRAAMVNQGSPMGDPRVSMQMQRRQ
330 340 350 360 370 380 390

MMMARRRMMAGKHEAPRTAFVGRLRQTMQKKK
400 410 420 422

TRINITY_DN27824_c0_g1_i4_ GMP-rich protein.prot (121 aa)

MMGMMNPYMMGMGGLGMMGMMNPYMMGMGGLGMMGMMNPYMMGMGGLGMMGMMNPYMMGMGGLGM
1 10 20 30 40 50 60

MGMMNPYMMGMGGLGMMGMMNPYMMGMGGLGMMGGYGMMGGLGMMGGLGMMTGYKK
70 80 90 100 110 120 121

**Fig. S5-3 RLDC and GM-rich protein sequences in cluster 4 and 8 marker genes**

**Fig. S5 RLDC and GM-rich protein sequences in cluster 4 and 8 marker genes**

The amino acid sequences for RLDC and GM-rich genes which are cluster 4 or 8 marker genes were shown. The visualization was conducted by SnapGene viewer (https://www.snapgene.com/snapgene-viewer). Low complexity regions, as detected by NCBI Conserved Domain search function (https://www.ncbi.nlm.nih.gov/Structure/cdd/wrpsb.cgi), are highlighted in red.

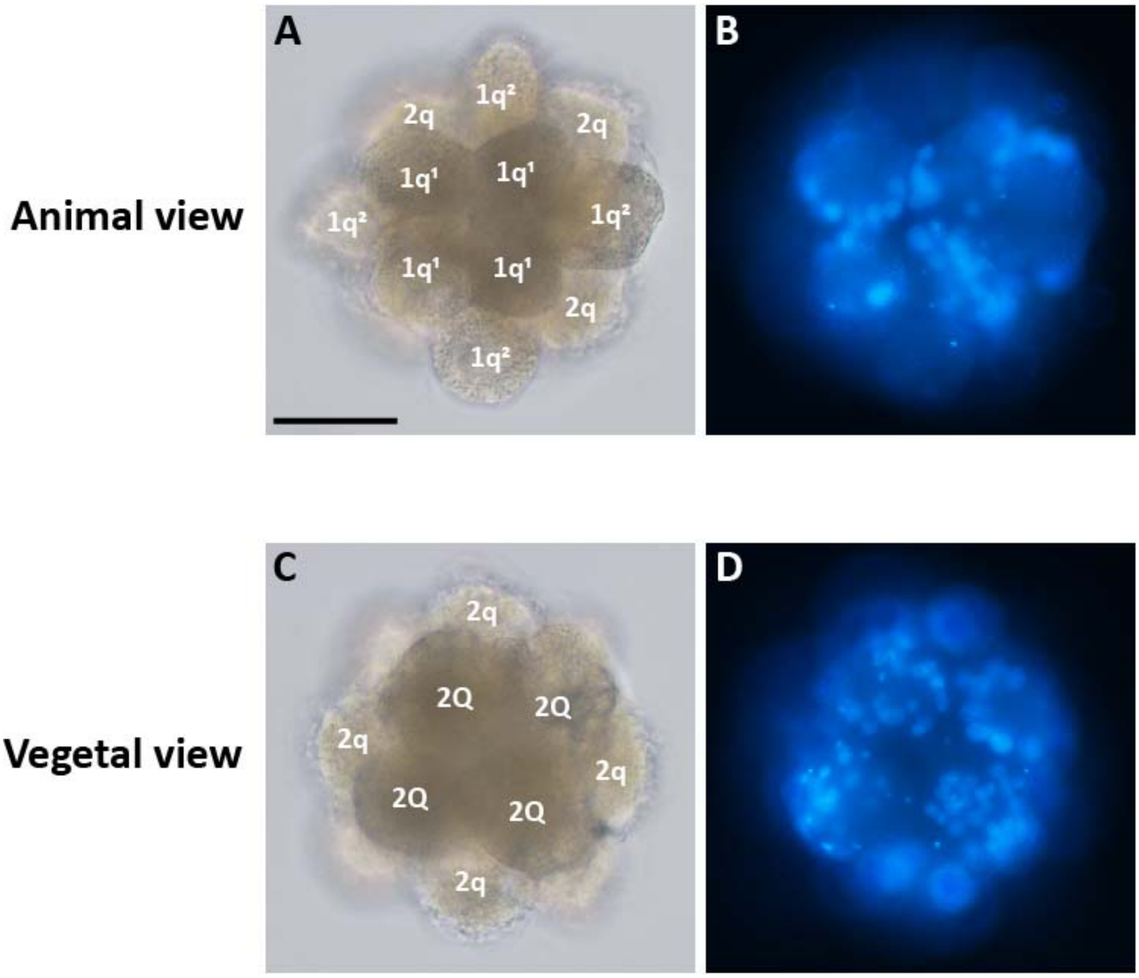


**Figure S6 The 9 hpf embryos blocked in cytokinesis from the 16-cell stage**

(A-B) The animal view of the cytokinesis-arrested embryo imaged using bright field microscopy (A) and fluorescence microscopy with DAPI-stained nuclei (B). (C-D) The vegetal view of the cytokinesis-arrested embryo imaged by bright field microscopy (C) and fluorescence microscopy with DAPI-stained nuclei (D). Scale bar = 50 µm.

| Gene name | Forward | Reverse + T3 sequence (underlined) |
|---|---|---|
| ***Hox1*** | CCCCACGGATTGGGGAATGGGGATATATTA | ATTAACCCTCACTAAAGGGACTTTTGTTTCATTCGTCGAT |
| ***Hox4*** | TCATGAGTTCGTTTTTGATGAACCCTGGAA | ATTAACCCTCACTAAAGGGAATTCTGCGATTTTGAAACCA |
| ***Mist*** | CACCAGACTTCAATTCCAGAAAGCGAGGAG | ATTAACCCTCACTAAAGGGATTTCCGGATCAGGAGCTTGG |
| ***Tyr*** | AAGAGGTCGAATGAATCCAAAGGCCGAACC | ATTAACCCTCACTAAAGGGAACTGGTGGCTGTAATCTTGG |
| ***Dpp*** | CGTTGAACCTGAGGAAGCATGGGAATGAGA | ATTAACCCTCACTAAAGGGAACAACAGGCTTTTGGTACGG |
| ***CS1*** | ACCAGATTTATCGGCCATCCCTGCAACAAC | ATTAACCCTCACTAAAGGGAGGTGGGCCAAGTGATATCTG |
| ***Tektin*** | TCAAGGAGACCGACACCCGAACTAGAAATC | ATTAACCCTCACTAAAGGGACCTTTTCTCATTCCCATACA |
| ***LR74*** | CAAGCCACAGGTGGTTCCTCAAAATAAAGT | ATTAACCCTCACTAAAGGGAAACGCTTTGAACCTCTTTCA |
| ***Unchar-2260*** | CTTCAACATGGCTGTGGGAAAATGAGTTTA | ATTAACCCTCACTAAAGGGATTCTTTAATTGCAGGTACGC |
| ***Sp6-9*** | CAATGTTAGGGGATCCACAGTCATATGGAG | ATTAACCCTCACTAAAGGGAGGCTGGTGATCCTGTACTAT |
| ***Dlx*** | GTAAATCCGCTTTTATGGAGCTACAACAAC | ATTAACCCTCACTAAAGGGAGTTGATACCAGGAATAAT |
| ***FoxAB*** | GCGTCACCAACACTTGATGGTTCTCAACTT | ATTAACCCTCACTAAAGGGAAAAGAGGAACTCAGCCCATT |
| ***Erg*** | CCACGACTGGGAACCAGTTGAATAACTTCC | ATTAACCCTCACTAAAGGGAATGGCCAATACGGACTCGTT |
| ***Alx*** | ATTAGGCACCAAACGACACCCAGACATGGA | ATTAACCCTCACTAAAGGGAACGCTGCTAGCTTGAACTGC |
| ***T23O*** | ATTCCTTATCGATGACGTCTGCAGTTCACG | ATTAACCCTCACTAAAGGGACACCATTGATTAAATCTGCT |
| ***KMO*** | AATGAGACCAGATATTCGCACACAACTTCA | ATTAACCCTCACTAAAGGGAAACAACAATACCAACAACCA |
| ***Blimp1*** | CGTGTTTGTCCAAATAGAGCTCAAGCATCG | ATTAACCCTCACTAAAGGGAGTGGTGATGAAGGGAAGATT |

**Table S1** Primers used for gene isolation
